# Genetic Diversity, Taxonomic Insights, and Geographic Distribution of *Bubo bubo* Subspecies in Iran

**DOI:** 10.1101/2025.07.27.667027

**Authors:** Arya Shafaeipour, Elham Rezazadeh, Behzad Fathinia, Michael Wink, Keramat Hafezi, Urban Olsson

**Affiliations:** Department of Biology, Yasouj University, Yasuj, Iran; Department of Animal Biology, Faculty of Biological Science, Kharazmi University, Tehran, Iran; Institute of Pharmacy and Molecular Biotechnology, Heidelberg University, Im Neuenheimer Feld 364, 69120, Heidelberg, Germany; Jondi Shapour University, Ahwaz, Iran; Dept of Biology and Environmental Sciences, University of Gothenburg, Göteborg, Sweden

**Keywords:** *Bubo bubo*, Iran, mitochondrial DNA, cytochrome b, genetic diversity, phylogenetic analysis, population expansion

## Abstract

The Eurasian Eagle Owl (*Bubo bubo*) exhibits a complex taxonomic structure with multiple subspecies across its broad geographic range, including southwestern Asia. While much is known about the species’ distribution in Europe and Asia, the taxonomic status and distribution of *B. bubo* subspecies in Iran remain poorly understood. This study aims to explore the genetic diversity and taxonomic relationships of *B. bubo* subspecies in Iran through mitochondrial gene analysis, focusing on nucleotide sequences of the cytochrome b (cytb) gene. A total of 36 samples collected from diverse localities across Iran, representing different ecological regions. Phylogenetic analysis revealed two distinct clades: one comprising western Iranian samples clustering with European (mostly from Germany) samples, and the other consisting of eastern and southern Iranian samples that shared haplotypes with populations from China. Notably, the western Iranian clade exhibited a minimal genetic distance from European populations, while the eastern Iranian clade was genetically similar to Chinese *B. bubo* subspecies, including *B. bubo turkomanus*. The genetic diversity indices for the Iranian population were relatively low, with three identified haplotypes and low nucleotide and haplotype diversity. Demographic analyses indicated a potential population expansion for the species, supported by a unimodal mismatch distribution and negative neutrality indices, although these results were not statistically significant for the Iranian population. The findings suggest that two subspecies, *B. bubo nikolskii* and *B. bubo interpositus*, are likely present in Iran, aligning with recent taxonomic assessments, but further investigation with additional genetic markers is needed to clarify the deeper evolutionary relationships among these populations. This study contributes new insights into the genetic landscape of *B. bubo* in Iran and the broader Palearctic region, with implications for conservation and future taxonomic revisions.

**Simple Summary:** This study examines the genetic diversity, taxonomic relationships, and geographic distribution of *Bubo bubo* subspecies in Iran, utilizing mitochondrial gene analysis, with a particular focus on the cytochrome b (cytb) gene. The research, based on 36 samples from various regions in Iran, reveals two distinct clades: one aligning with European populations and the other sharing haplotypes with Chinese subspecies. While genetic diversity in Iranian populations was relatively low, the findings highlight the presence of *B. bubo nikolskii* and *B. bubo interpositus* subspecies in Iran, emphasizing the importance of further genetic investigations for conservation and taxonomic revisions in the Palearctic region.

## 1. Introduction

Owls (Strigiformes) are among of the most recognizable birds, and their specialized traits, such as advanced auditory capabilities [1, 2] and silent flight [3, 4] support their lifestyle as nocturnal predators. These adaptations have allowed over 256 species of owls [5-7] to inhabit a wide variety of habitats worldwide, from tundra to dense forests [8]. Among these predators, the Eurasian Eagle Owl (*Bubo bubo*) stands out as one of the largest representatives of Strigiformes. The species is distributed troughout the Palearctic, with 16 subspecies found in various regions, experiencing significant declines in recent decades. Althoug these declines are well documented in some areas, literature indicates that our understanding of the species’ status in southwestern Asia remains limited [9].

The taxonomic classification of *Bubo bubo* has been a topic of ongoing debate, particularly due to its diverse geographic distribution, variations in plumage and differences in size [10, 11]. This complexity has led to the recognition of multiple subspecies across the range of this species. For instance, Wink and Heidrich [12] identified 16 subspecies, while Gill [13] recognized up to 17 subspecies. Historically, two subspecies have been reported in certain parts of Africa, including *B. b. interpositus* and *B. b. ascalaphus* [10, 11] with the latter, the Pharaoh eagle-owl (*Bubo ascalaphus*), elevated to a specific rank thanks to advancements in molecular and ecological approaches in taxonomic studies. Thus, *B. b. interpositus* is the only representative of Eurasian eagle-owl inhabiting northeastern regions of this continent [14-16]. Similiar to Africa, the Indian subcontinent also hosts two subspecies of the Eurasian eagle-owl: the Rock Eagle Owl, *B. b. bengalensis*, which occupies peninsular India, and the Himalayan Eagle-Owl, *B. b. hemachalana*, found in the northwestern Himalayas [17]. The Rock Eagle Owl was elevated to a specific rank, while the Himalayan Eagle-Owl was suggested to be synonymous with *B. b. turcomanus* [15, 18]. More recent studies, such as those by Meng et al., [19], proposed the existence of 12 subspecies, while Clements et al., [20] suggested a total of 16 subspecies, including *B. bubo nikolskii* and *B. bubo interpositus*, both found in Iran.

The Middle East, particularly Iran, is crucial for understanding the distribution of these subspecies [21]. The diverse landscapes of Iran, characterized by arid deserts and mountainous regions, likely provide various habitats that could support multiple *Bubo bubo* subspecies [22]. It has been proposed that the region’s topographical complexity may have offered stable habitats, especially for vertebrates, during historical ice ages [23, 24]. Currently, Iran represents the southernmost limit of the species’ distribution, with known populations primarily located in the eastern half of the country, along with a few isolated occurrences. However, comprehensive studies focusing on the specific distribution patterns and taxonomic affiliations of *B. bubo* subspecies within Iran are still lacking [25].

To address this knowledge gap, we analyzed the nucleotide sequences of a mitochondrial gene (cytochrome b) to assess the genetic diversity and population structure of *B. bubo* subspecies sampled from various localities throughout Iran, and for comparison, from Europe. This study aims to enhance our understanding of the genetic variation and demographic processes affecting this species within the Iranian landscape.

## 2. Materials and Methods

In total, 36 samples (crashed, shot, or sick birds), including blood (22), muscle (10), and feather (4) were obtained from different locations in Iran (Figure 1). All tissue samples were stored at – 20°C or in 96% ethanol. DNA was extracted using the Sinuhe Biotechnology Kit (Sinuhebiotech, Iran), according to the manufacturer’s instructions. We sequenced the mitochondrial cytochrome b (cytb) gene for 36 samples. Amplification and sequencing followed the protocols described in Olsson et al., [26]. We amplified a fragment of the mitochondrial cytochrome b (cytb) gene using the primers ND5-Syl (5’-GGCCTAATCAARGCCTACYTAGG-3’) and mtF-NP (5’-GYTTACAAGACCAATGTTT-3’) [27]. PCR amplifications were performed in a reaction volume of 25.4 µl, containing 12 µl ddH2O, 10 Mastermix, 0.7 µl of each primer, and 2 µl template DNA. The thermocycling conditions for amplifying cytochrome b were outlined as follows: initial denaturation at 95 °C for 5 minutes; 40 cycles of denaturation at 95 °C for 45 seconds, annealing at 50 °C for 45 seconds, and extension at 72 °C for 2 minutes; followed by a final elongation at 72 °C for 5 minutes.

**Figure 1.**
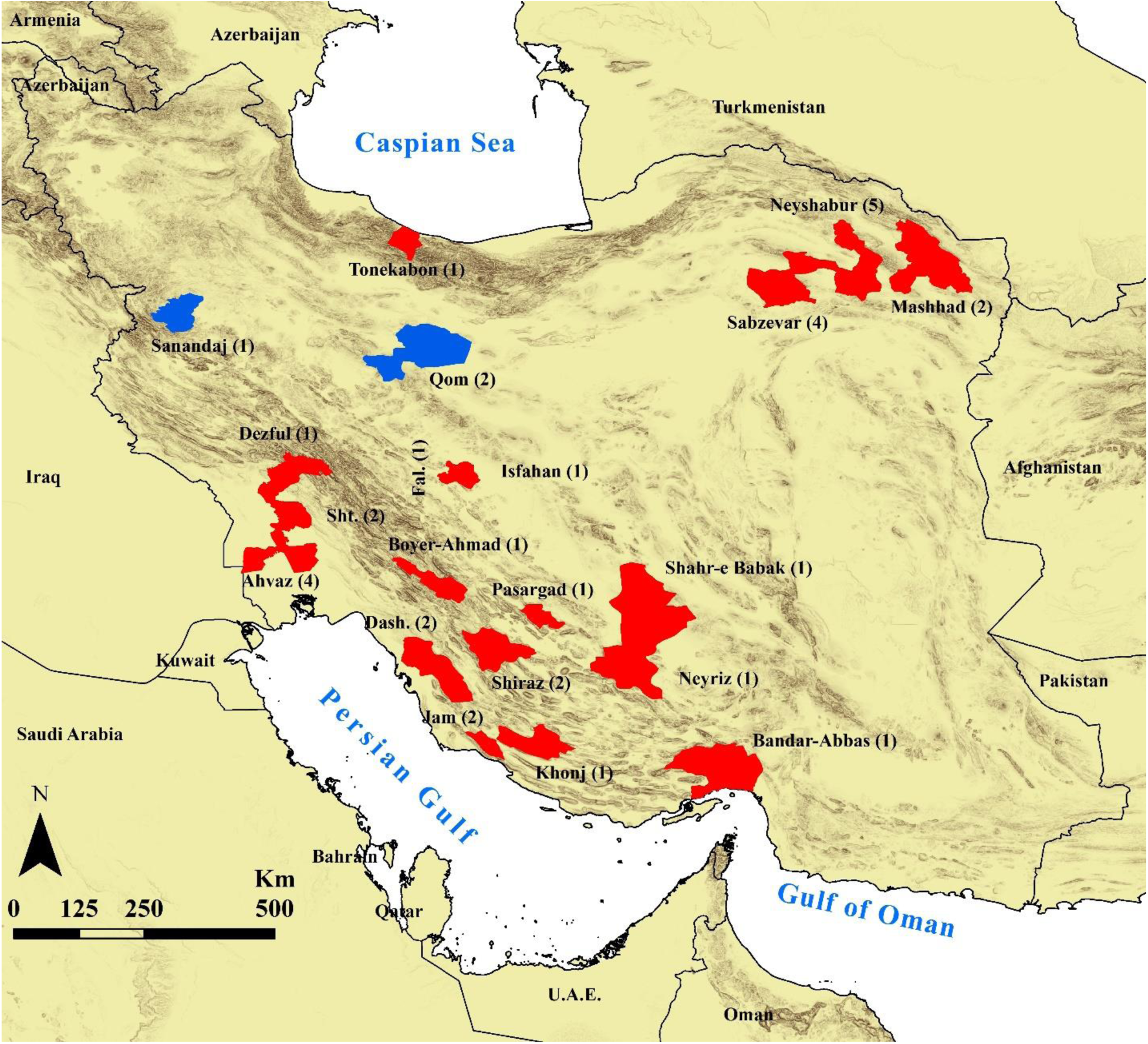
Map of sampled localities of *Bubo bubo* in Iran. Blue and red polygons indicate the locations of samples assigned to western and eastern/southern Iran. Values within the parentheses indicate the number of individuals collected from each locality. Abbreviations: Dash. =Dashtestan; Sht. = Shushtar; Fal. = Falavarjan.

Additional sequences from GenBank were also included for a more comprehensive molecular dataset (Supplementary Table S1). Sequences were aligned by the clustal W algorithm [28] through Bioedit software [29]. The degree of substitution saturation to identify if the dataset is reliable for phylogenetic analyses was determined through comparison of ISS vs. ISS.c in DAMBE v. 5.3.109 [30]. Pairwise genetic distances among ingroup-only sequences were computed with MEGA 12 [31]. The best model for genetic distance in the ingroup dataset was calculated in MEGA 12 [31], resulting in the HKY model. As this model is not installed in MEGA’s distance options, we used the closest alternative, the Tamura-Nei model (TN93). Details on specimens, GenBank accession numbers, and corresponding localities can be found in Table S-1.

Phylogenetic analyses utilized sequence data derived from an amplified fragment of mitochondrial DNA (mtDNA) genes. The appropriate models of evolution were determined through PartitionFinder 1.0.1 for Bayesian Inference and Maximum Likelihood analyses [32]. Bayesian Inference (BI) and Maximum Likelihood analyses were conducted for Cytb using MrBayes v.3.1 [33] and RaxMLGUI [34], respectively. Bayesian inference was simulated under HKY+I model for 2×10^6^ generations, and log likelihood values of sample points were plotted against generation time. The adequacy of BI analysis was assessed through the “Average standard deviation of split frequencies”, “ESS,” and “PSRF”. A 25% burn-in discarded initial trees, and the remaining trees were amalgamated to form a 50% majority consensus tree. Nodal robustness of the BI trees were assessed through Bayesian posterior probabilities, considering BPP> 0.90 as indicative of strong support. Maximum likelihood analysis was simulated under GTR+G model using the following parameters: a random starting tree, 100 replicates, and assessing nodal support with 1,000 bootstrap pseudoreplications [35].

To reveal the phylogenetic relationships, the TCS haplotype network was constructed using PopART V. 1.7 [36]. DnaSP V. 5.10 [37] was employed to calculate parsimony informative sites and total mutations.

### 2.1. Demography

Demographic analyses were conducted on the mtDNA marker Cytb, utilizing the complete dataset. The number of polymorphic sites, nucleotide diversity, average nucleotide diversity, number of haplotypes, and haplotype diversity were calculated using DnaSP v5.10 [37]. Pairwise FST values among different taxa were assessed with Arlequin v3.5 [38], and 1,000 simulated samples were used to evaluate any significant differences (p < .05).

Additionally, two commonly used methods were utilized to explore the demographic history of the populations. First, mismatch distributions for mtDNA marker were generated separately. A unimodal distribution, indicative of a sudden population expansion, was compared to a multimodal distribution, suggesting demographic equilibrium. This analysis was performed using Arlequin with 1,000 simulated random samples. Second, demographic equilibrium for each phylogroup was evaluated through Fu’s Fs [39], sum of squared deviation (SSD), raggedness index (rg) [40], and Tajima’s D [41], with significance assessed using 1,000 simulated samples in Arlequin. The Ramos-Onsins and Rozas test R2 [42] was estimated using 10,000 coalescent simulations in DnaSP. A significance threshold of p <.05 was applied to all statistics, except for Fu’s Fs, where p < .02 was deemed significant [38].

## 3. Results

The following results obtained from 94 sequences, including 1 sequence of *Bubo poensis* as outgroup and 93 of *B. bubo* as ingroup, ranging from Europe through the Middle-East to China. Nucleotide frequency distribution in the partial Cytb gene (584 bp) includes: A, 25.81%; C, 35.03%; T, 23.88%; and G, 15.28%. A total of 54 variable and 19 parsimony-informative sites were obtained for the ingroup dataset, while they were 106 and 28 sites in the ingroup and outgroup dataset combined (Table S1). There is a low substitution saturation (ISS = 0.01; ISS.c = 0.703; P<0.0005) in the Cytb dataset, indicating the sequences’ suitability for phylogenetic analyses. Notably, a Western European cluster can be distinguished from an Eastern Asian cluster. The two approaches (ML and BI inferences) revealed that samples from Iran clustered into one of two distinct clades. In other words, samples from western Iran are nested within the European samples, while those from southern and eastern Iran are clustered with samples from China (Figures 2-3). The TN93 genetic distances among different clades are very low, indicating little genetic divergence within this species (Table 1) and between eastern and western clades, being 0.007. Network analysis using the TCS method corroborated the phylogenetic analyses, identifying two clusters separating samples from eastern and western populations, albeit differing by only a few base substitutions. Almost all samples from western and eastern Iran were identical. Only the two singletons from eastern (Hap_14) and southern (Hap_13) Iran differed by one mutation from the eastern majority haplotype (i.e. Hap_12), which had divided into two different haplotypes (Figure 4, Table 2).

**Figure 2.**
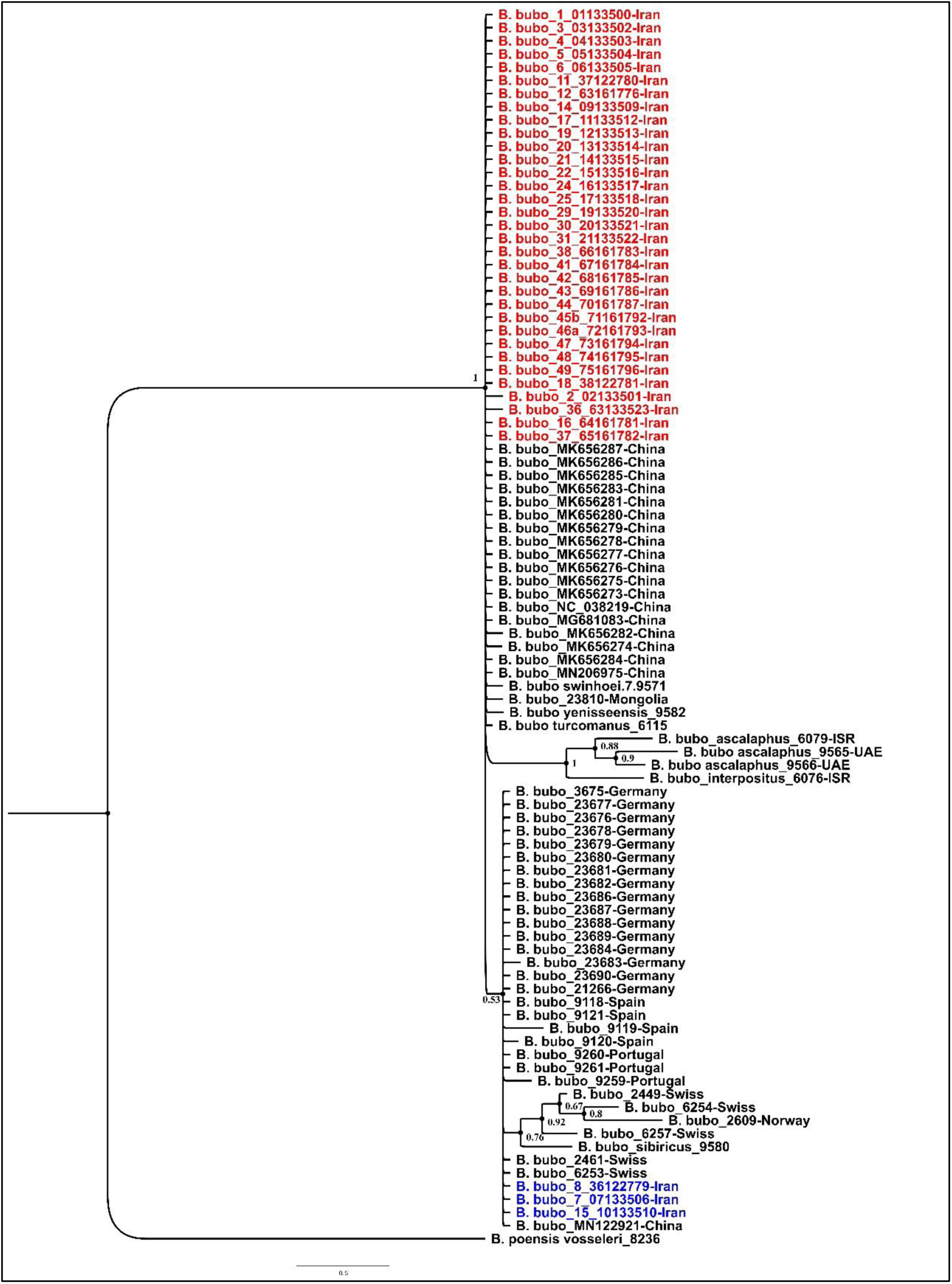
The Bayesian inference tree shows the relationships among populations of *Bubo bubo* resulting from 94 partial sequences of the mitochondrial Cytb gene. Posterior probability values greater than 0.5 are

**Figure 3.**
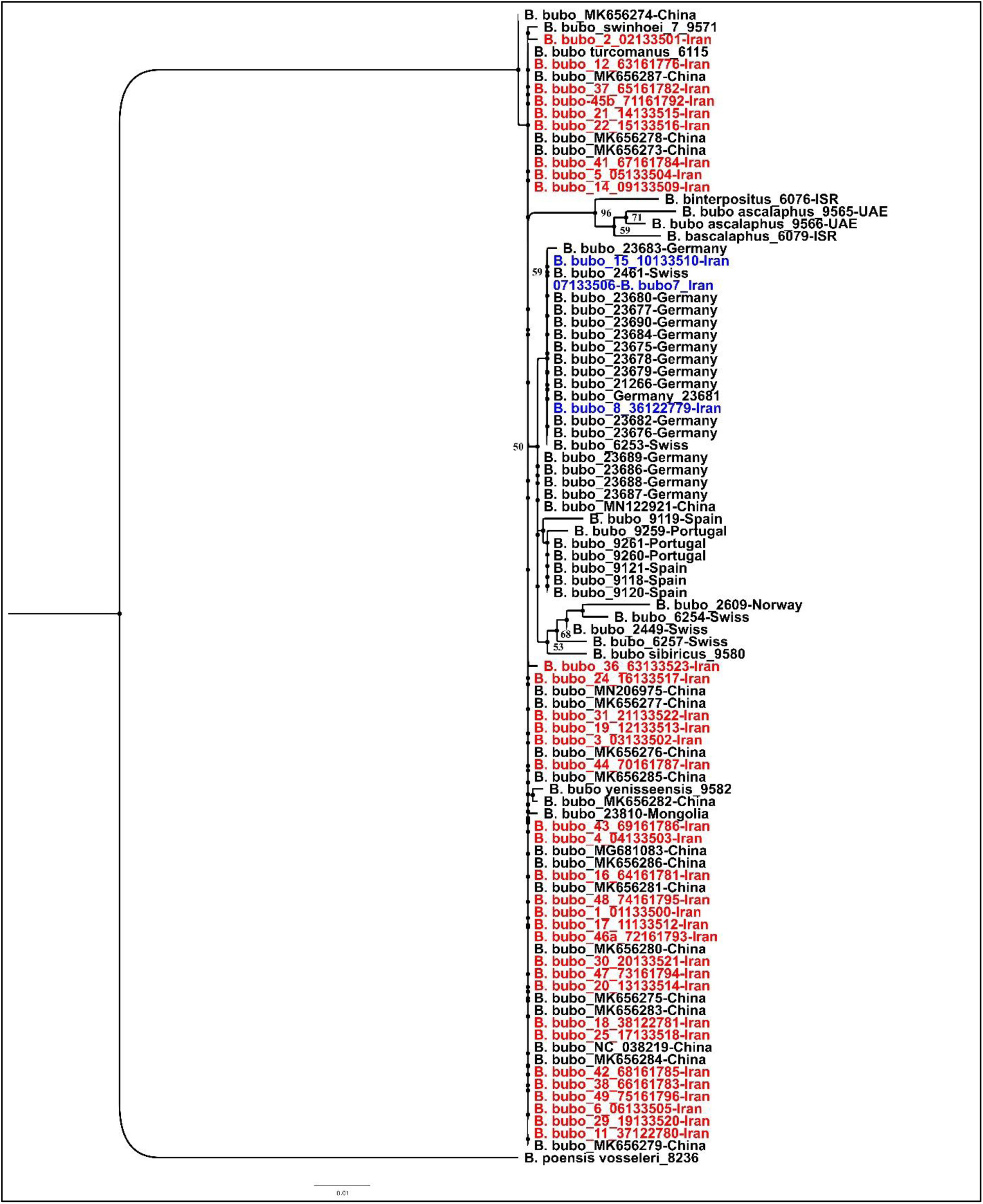
The maximum Likelihood tree shows the relationships among populations of *Bubo bubo* resulting from 94 partial sequences of the mitochondrial Cytb gene. Bootstrap supporting values greater than 50 are

**Figure 4.**
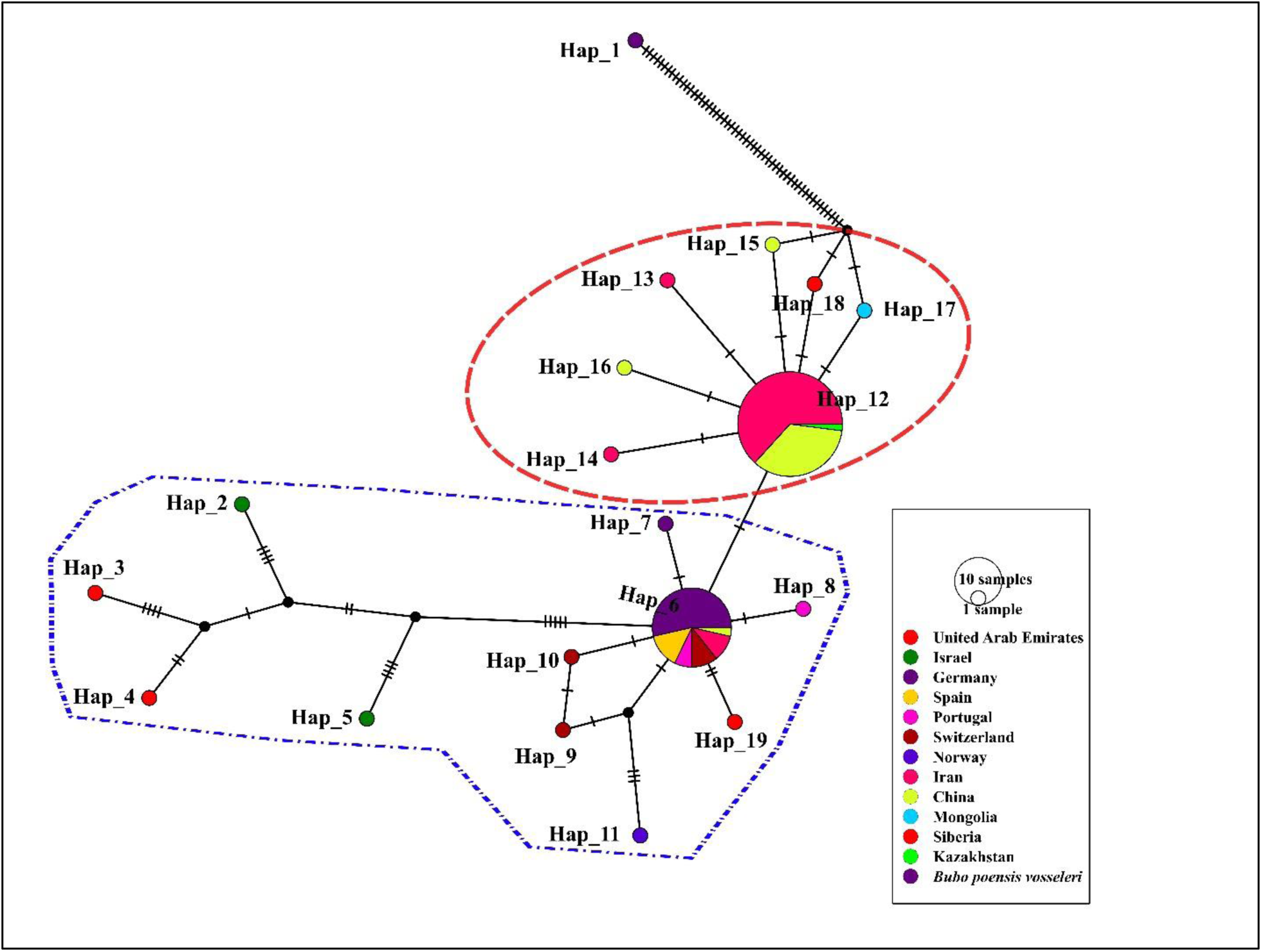
TCS networks of *Bubo bubo* populations for the Cytb marker. Each circle represents a haplotype, and the size of the circle is proportional to that haplotype’s frequency. The hatch marks at each branch indicate the number of nucleotide exchanges. The red dashed ellipse and the blue dashed polygon indicate eastern and western clades, respectively.

**Table 1.** Genetic mean distance among 5 clades of *Bubo bubo* based on the Tamura-Nei model (values below diagonal) in the *Cytb* sequences. Standard error estimates are shown above the diagonal. As the table implies, the western Iran clade is closer to European population, while the eastern/southern clade is closer to the Chinese population.

| Clade | West Iran | E. & S. Iran | Europe | China | UAE+Israel |
| --- | --- | --- | --- | --- | --- |
| West Iran |  | 0.003 | 0.001 | 0.003 | 0.006 |
| E. & S. Iran | 0.004 |  | 0.002 | 0.000 | 0.005 |
| Europe | 0.003 | 0.005 |  | 0.002 | 0.005 |
| China | 0.004 | 0.001 | 0.005 |  | 0.005 |
| UAE+Israel | 0.025 | 0.022 | 0.026 | 0.022 |  |

**Table 2.**
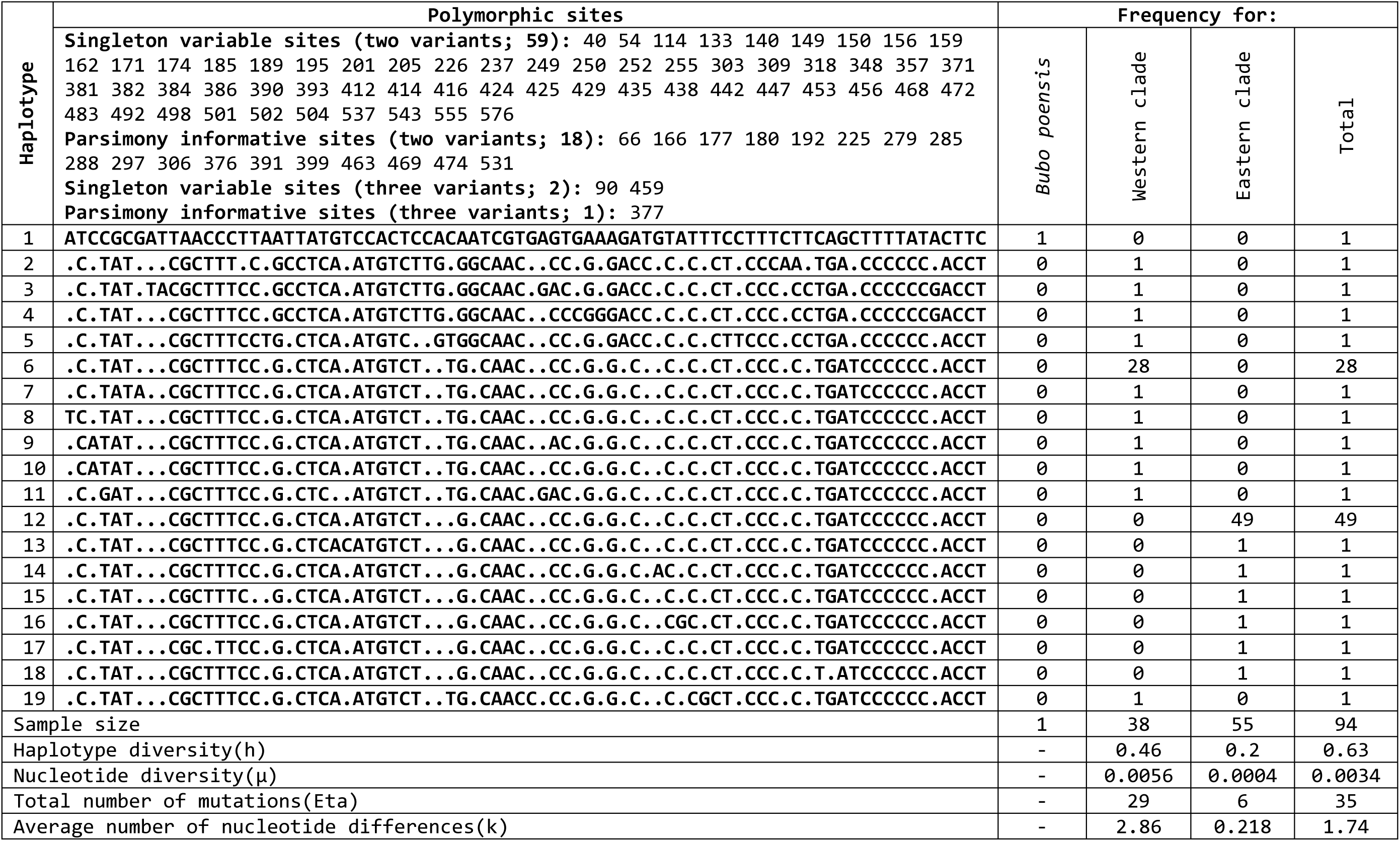
The comparison of different parameters of the haplotypes of the Cytb gene within and between populations of *Bubo bubo*, used in this study.

Analyses of different parameters of genetic diversity in *Bubo bubo* for the Iranian populations, along with the complete dataset revealed a total of 4 and 18 haplotypes, respectively. Both nucleotide and haplotype diversities are low in the Iranian population and in the complete dataset (Tables 2-3).

**Table 3.** Summary of genetic diversity of the members of *Bubo bubo* samples based on Cytb. The significance of calculated statistics is indicated by asterisks in the parentheses (*p<.05, **p<.02). Abbreviations: Sample size (No.); number of haplotypes (H); haplotype diversity (Hd); standard deviation (SD); nucleotide diversity (π); number of segregating sites (S); average number of differences (k).

| Population | No. | H | Hd±SD | $\pi$±SD | S | K | Tajima D | Fu's FS | R2 |
| --- | --- | --- | --- | --- | --- | --- | --- | --- | --- |
| **Total** | 93 | 18 | 0.63±0.04 | 0.003±0.0008 | 35 | 1.74 | -2.32 (<0.01**) | -9.55 (<0.02**) | 0.025 |
| **Iran** | 36 | 4 | 0.26±0.09 | 0.0007±0.0002 | 4 | 0.42 | -1.37 (>0.1) | -1.48 (>0.1) | 0.07 |

For the Iranian population, non-significant values for Fu’s Fs, Tajima’s D, and R^2^ suggested no evidence of recent population expansion (Table 3), as reflected by the unimodal distribution profile in the mismatch distribution (Figure 5). Conversely, Tajima’s D and Fu’s Fs indices were significantly negative for the complete dataset. These negative indices, along with the L-shaped mismatch distribution, may indicate a recent population expansion following a bottleneck event (Figure 5).

**Figure 5.**
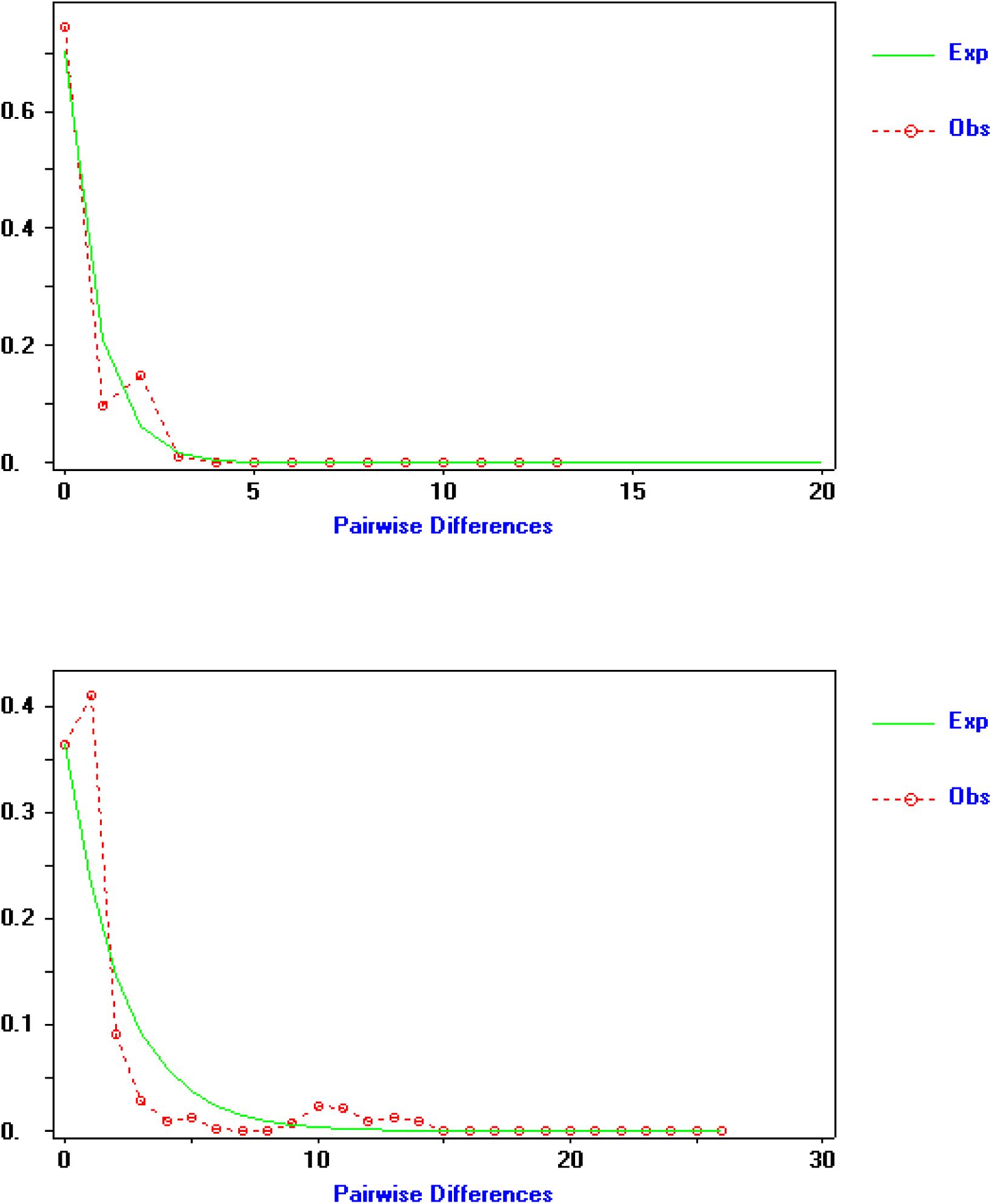
Mismatch distribution inferred from Cytb marker in *Bubo bubo* for the Iranian populations (above) and across the species (Below).

## 4. Discussion

This study provides valuable insights into the genetic structure and diversity of *Bubo bubo* populations in Iran, revealing distinct genetic patterns within the region. Our findings suggest two primary clades in the Iranian population: western and eastern/southern groups, which exhibit low genetic divergence but distinct genetic profiles. These findings should be interpreted in the context of broader Eurasian studies, particularly those investigating *B. bubo* populations in Europe, China, and some parts of Asia. Hereafter, the current results are compared with previous findings, focusing on genetic diversity, population structure, and their implications for subspecies identification and demographics.

### 4.1. Genetic Structure

Our Bayesian phylogenetic analysis and median-joining network indicate that *B. bubo* populations in Iran form two main clades, with the western Iranian populations clustering closely with European samples, particularly those from Germany. The eastern and southern Iranian populations showed a close genetic similarity to samples from China, suggesting a possible genetic link across these regions. However, the phylogenetic support for these clusters was weak, as indicated by low posterior probabilities. These clades are also supported by a few unique base substitutions, which attenuate the results of this study. The genetic distance between eastern and western populations was relatively small, with a maximum TN93 genetic distance of 0.007, and the network analysis revealed only two base substitutions separating these clades. This suggests that while there is some differentiation between eastern and western populations of Iran, their genetic divergence is minimal, which could imply ongoing gene flow.

In comparison, a study in China based on analyses conducted on complete mitogenome has identified three distinct subspecies of *B. bubo*: *B. b. kiautschensis*, *B. b. ussuriensis*, and *B. b. tibetanus* [19]. These subspecies exhibit clear genetic distinctions, characterized by low nucleotide but high haplotype diversity observed within each subspecies. The phylogenetic relationships of these subspecies align with geographic and environmental gradients across China. In contrast, our study showed low genetic diversity within the Iranian population, with only four haplotypes identified from 36 samples, and weak phylogenetic support for clustering with European or Chinese populations. This suggests that Iranian populations may represent a genetic admixture of European and Asian lineages, reflecting their geographic position at the intersection of multiple migration routes.

### 4.2. Genetic Diversity and Neutrality Tests

Our results showed low genetic diversity in the Iranian populations, with a total of only four haplotypes identified from 36 samples. Both nucleotide and haplotype diversity were low, and neutrality tests (Tajima’s D, Fu’s FS, and R2) were non-significant for the Iranian populations, indicating no evidence of recent population expansion. However, this finding was not supported by the mismatch distribution, which suggested an expansion pattern. This discrepancy may reflect the local genetic landscape, where demographic changes could be influenced by regional ecological factors or historical population bottlenecks.

In contrast, studies in other regions have reported evidence of population expansion in *B. bubo* populations. For instance, a study on the Iberian population found a star-like phylogenetic pattern and evidence of recent expansion in northern Europe, but not in the Iberian population [43]. Similarly, studies in China have identified multiple subspecies with distinct genetic patterns, suggesting historical population expansions and contractions influenced by climatic and geographic factors [19]. These differences highlight the complex demographic history of *B. bubo* across Eurasia and underscore the need for region-specific conservation strategies. Isolation by distance (IBD) can be another explanation for the minimal differences observed between the two clades. Subtle genetic differences can be a result of Isolation by distance (IBD) rather than population structure. Indeed, IBD is occurred when genetic relatedness among individuals decreases as geographical distance increases, especially in those organisms having limited dispersal capabilities. Conversely, IBD can even shape genetic structure of distant populations [44].

### 4.3. Subspecies Identification

One of the key goals of this study was to assess the subspecies identity of Iranian *B. bubo* populations. A problematic topic in subspecies delineation occurs when the used dataset does not support population structure. This problem might be occurred when mall datasets (e.g., limited loci or samples) lack power to detect subtle structure, or historical subspecies designations, especially those based on morphology does not concordant with genetic patterns [45]. Recent checklists of birds suggest that two subspecies of *B. bubo* are present in Iran: *B. b. nikolskii* in eastern regions and *B. b. interpositus* in western regions [17]. Our phylogenetic results indicate that the western Iranian population is genetically similar to European populations, particularly those from Germany, which challenges the previous assumption that it belongs to *B. b. interpositus*. Furthermore, the eastern and southern Iranian populations shared genetic similarities with Chinese populations, with some haplotypes overlapping with those from *B. b. kiautschensis* or *B. b. turkomanus*. This raises the possibility that Iranian *B. bubo* populations could be assigned to *B. b. turkomanus*, although further research using additional markers or complete mitochondrial genomes would be needed to confirm this hypothesis.

### 4.4. Population Dynamics and Climate History

The demographic patterns observed in our study also offer insights into the population dynamics of *B. bubo* in Iran. The non-significant neutrality test values suggest a stable or slowly fluctuating population, rather than one that has recently expanded. This contrasts with the unimodal distribution and negative neutrality test values found for the broader dataset of *B. bubo* populations, which indicates recent expansion across Eurasia. These findings align with previous studies suggesting regional differences in population dynamics in *B. bubo*. For example, studies have found evidence for demographic stability in certain Mediterranean populations of *B. bubo*, likely due to stable climatic conditions in these areas [46]. In contrast, populations in northern Europe and central Asia may have undergone population expansions during periods of favorable climatic conditions, such as the end of the last glacial maximum.

## 5. Conclusions

In summary, the findings from this study contribute significantly to our understanding of the genetic diversity and structure of *B. bubo* populations in Iran. While there is some genetic differentiation between Western and Eastern populations, their overall low divergence suggests ongoing gene flow, and the genetic diversity within Iran remains relatively low. Further research, incorporating additional molecular markers and comprehensive sampling across the species’ range, will be crucial for clarifying the subspecies status of Iranian populations and providing insights into their demographic history.

## Supplementary Materials

The following supporting information can be downloaded at: https://www.mdpi.com/article/?????/birds?????/s1. Table S1: Information on specimens used in this study, with voucher numbers, collecting localities, and GenBank accession numbers. YUZM: Yasouj University Zoological Museum.

## Authors’ contributions

All authors contributed to the study conception and design. Material preparation and data collection were performed by Arya Shafaeipour, Behzad Fathinia, Michael Wink, and Keramat Hafezi. Experimental procedures were conducted by Behzad Fathinia and Arya Shafaeipour. Analyses were performed by Behzad Fathinia, Elham Rezazadeh, Michael Wink, and Urban Olsson. The first draft of the manuscript was written by Arya Shafaeipour, Elham Rezazadeh, Behzad Fathinia, Michael Wink, and Urban Olsson, and all authors commented on previous versions of the manuscript. All authors read and approved the final manuscript. M. Wink thanks Hedwig Sauer-Gürth and Lona Wessels for carrying out the sequencing of the European samples.

## Institutional Review Board Statement

Not applicable.

## Data Availability Statement

Available from the first author on request.

## Acknowledgments

We thank members of the Bayqoosh Birdwatching Group in Iran for their assistance in obtaining samples.

---

**Supplementary Table S1.**
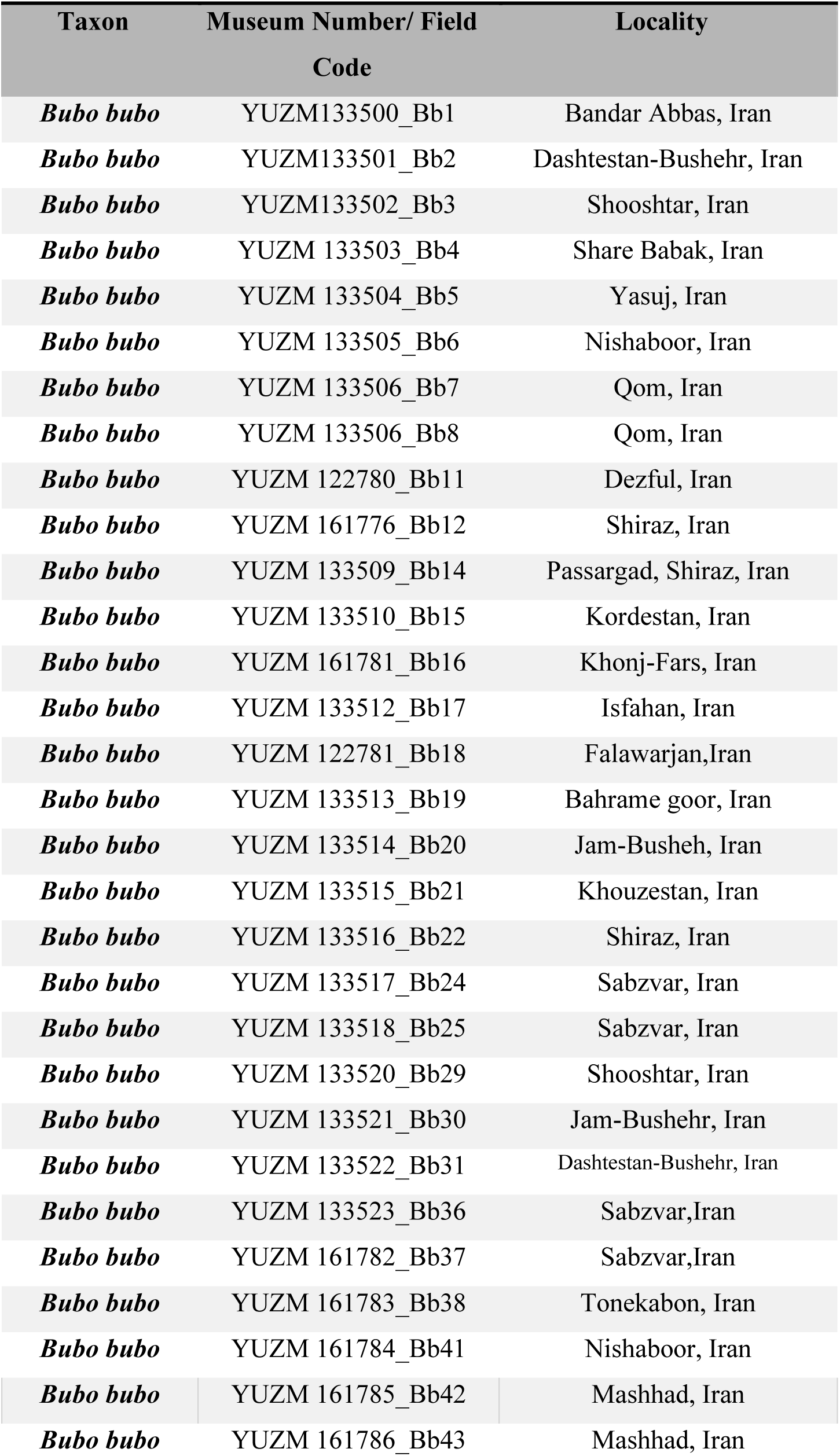
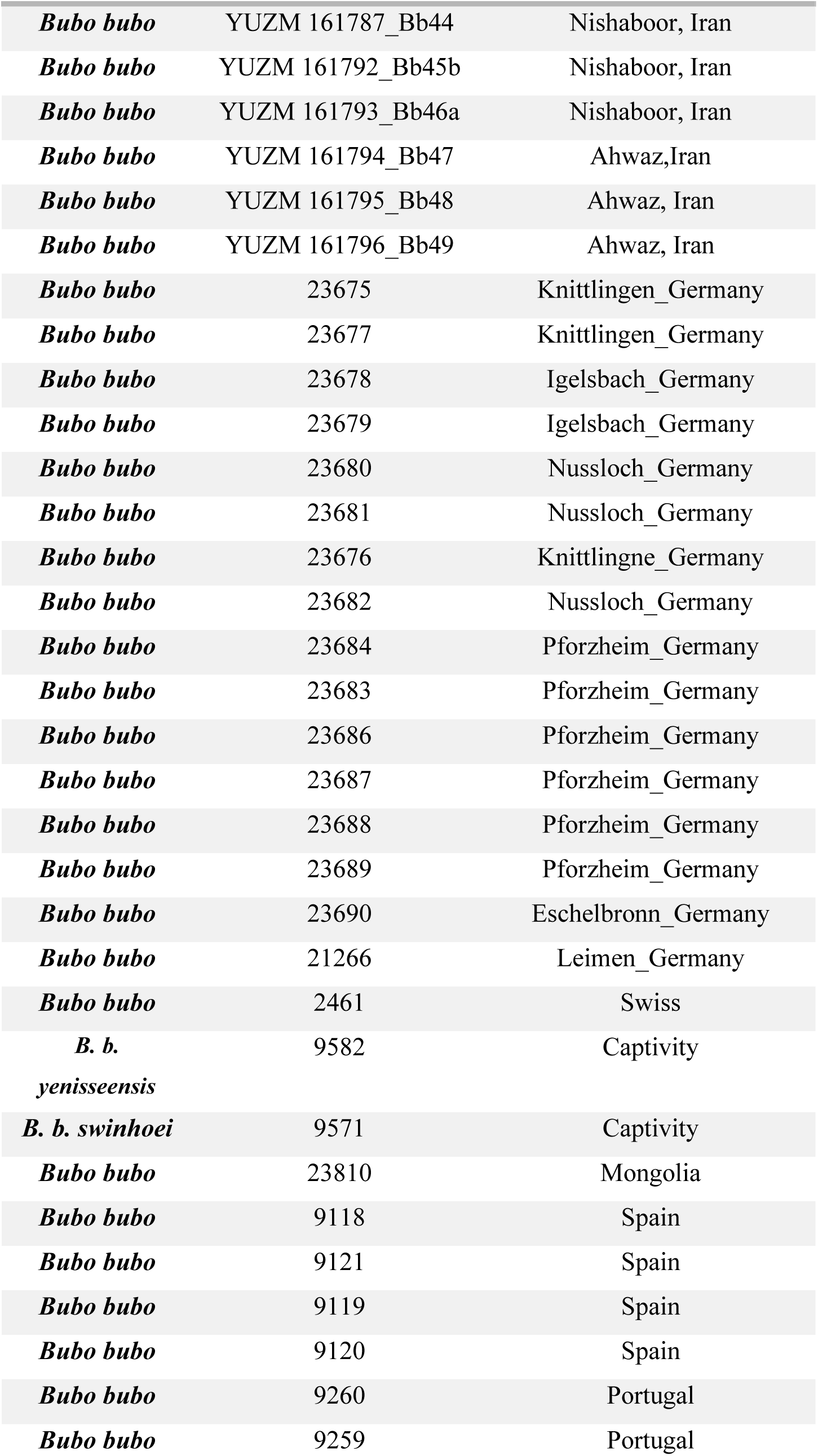
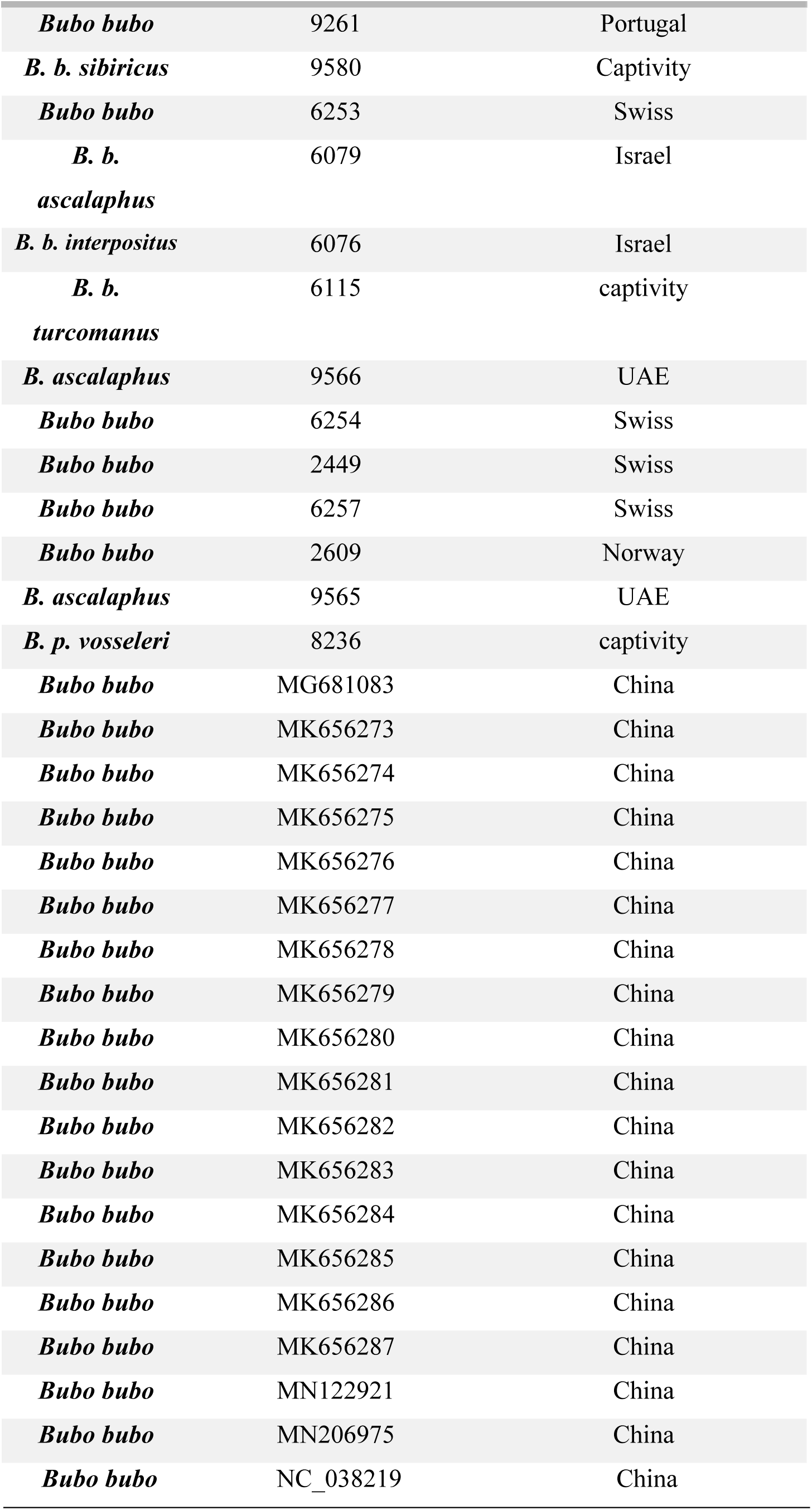
Information on specimens used in this study, with voucher numbers, collecting localities, and GenBank accession numbers. YUZM: Yasouj University Zoological Museum.

## References

1. Payne, R.S. Acoustic location of prey by barn owls (*Tyto alba*). J. Exp. Biol. 1971, 54(3), 535–573.

2. Konishi, M. How the owl tracks its prey: experiments with trained barn owls reveal how their acute sense of hearing enables them to catch prey in the dark. Am. Sci. 1973, 61(4), 414–424.

3. Sarradj, E.; C. Fritzsche; T. Geyer. Silent owl flight: bird flyover noise measurements. AIAA journal. 2011, 49(4), 769–779.

4. Geyer, T.; E. Sarradj; C. Fritzsche. Silent owl flight: comparative acoustic wind tunnel measurements on prepared wings. Acta Acust. united Ac. 2013, 99(1), 139–153.

5. Dickinson, E.; J. Remsen Jr; L. Christidis. The Howard & Moore Complete Checklist of the Birds of the World Eastbourne, UK: Aves Press, Eastbourne, 2013; Vol. Vol. 1, 2.

6. del Hoyo, J. All the Birds of the World: Lynx edicions, 2020.

7. del Hoyo, J.; N.J. Collar; D. Christie; A. Elliott; L. Fishpool; P. Boesman; G. Kirwan. Illustrated Checklist of the Birds of the World: Non-passerines: Lynx Ediciones, 2014.

8. Marks, J.; R. Cannings; H. Mikkola, J. del Hoyo, A. Elliott, J. Sargatal, D. Christie, E. de Juana. Typical owls (Strigidae). Handbook of the Birds of the World Alive (J. del Hoyo, A. Elliott, J. Sargatal, DA Christie, and E. de Juana, Editors). Lynx Edicions, Barcelona, Spain. 2018.

9. Salter, J.F.; C.H. Oliveros; P.A. Hosner; J.D. Manthey; M.B. Robbins; R.G. Moyle; R.T. Brumfield; B.C. Faircloth. Extensive paraphyly in the typical owl family (Strigidae). The Auk. 2020, 137(1), ukz070.

10. Hayashi, Y. Morphological Characters and the Source of Eagle Owl Bubo bubo Specimens in the Far East. J Yama. Ins. Ornitho. 1997, 29(1), 73–79.

11. Kim, H.-J.; H. Kim; S.-J. Park; C.-Y. Choi. Reversed Sexual Size Dimorphism and Morphological Sex Determination of the Smallest Subspecies of Eurasian Eagle-Owls (Bubo bubo kiautschensis). J. Raptor Res. 2024, 58(3), 329–337.

12. Wink, M.; P. Heidrich. Molecular evolution and systematics of owls (Strigiformes). Owls A Guide to the Owls of the World. 1999, 39–57.

13. Gill FB, W.M. Birds of the World. Princeton Univ.: Press, Princeton, 2006.

14. Sibley, C.G.; B.L. Monroe. Distribution and taxonomy of birds of the world. USA: Yale University Press, 1990; 1111.

15. Wink, M.; A.-A. El-Sayed; H. Sauer-Gürth; J. Gonzalez. Molecular phylogeny of owls (Strigiformes) inferred from DNA sequences of the mitochondrial cytochrome b and the nuclear RAG-1 gene. Ardea. 2009, 97(4), 581–591.

16. Wink, M.; H. Sauer-Gurth; I. Rogue. Molecular taxonomy and systematics of owls (Strigiformes)— An update. Airo. 2021, 29, 487–500.

17. Grimmett, R.; C. Inskipp; T. Inskipp. Birds of the Indian subcontinent IInd edn. Christopher Helm. India: Oxford University Press, 2011.

18. König, C.; W. Weick. Owls. A guide to the owls of the World ed.; S. edition. London: Christopher Helm, 2008.

19. Meng, M.; J. Ma; M.Y. Laghari; J. Ji. Genetic analysis of three wild Eurasian eagle-owl subspecies, B. b. kiautschensis, B. b. ussuriensis, and B. b. tibetanus, in Chinese populations. mtDNA Part B. 2020, 5(3), 3775–3777.

20. Clements, J.F.; P.C. Rasmussen; T.S. Schulenberg; M.J. Iliff; T.A. Fredericks; J.A. Gerbracht; D. Lepage; A. Spencer; S.M. Billerman; B.L. Sullivan; M. Smith; C.L. Wood. The eBird/Clements checklist of Birds of the World: v2024. 2024.

21. Khan, M.A., et al. Habitat diversity and Bubo bubo populations in Iran. J. Ornithol. Res. 2020, 18 (4), 310–325.

22. Adler, R.; S. Patel. Bubo bubo subspecies distribution in the Middle East. Wildlife Divers. Res. 2018, 7(4), 112–125.

23. Ahmadzadeh, F.; M. Flecks; M.A. Carretero; O. Mozaffari; W. Böhme; D.J. Harris; S. Freitas; D. Rödder. Cryptic speciation patterns in Iranian rock lizards uncovered by integrative taxonomy. PloS one. 2013, 8(12), e80563.

24. Rezazadeh, E.; M. Aliabadian; J. Darvish; F. Ahmadzadeh. Diversification and evolutionary history of brush-tailed mice, Calomyscidae (Rodentia), in southwestern Asia. Org. Divers. Evol. 2020, 20, 155–170.

25. White, L.M., & Garcia, R. A. Current status of Bubo bubo research in the Middle East. Ornitho. Progress. 2021, 14(3), 200–215.

26. Olsson, U.; P. Alström; P.G. Ericson; P. Sundberg. Non-monophyletic taxa and cryptic species— evidence from a molecular phylogeny of leaf-warblers (Phylloscopus, Aves). Mol. Phylogenetics Evol. 2005, 36(2), 261–276.

27. Stervander, M.; J.C. Illera; L. Kvist; P. Barbosa; N.P. Keehnen; P. Pruisscher; S. Bensch; B. Hansson. Disentangling the complex evolutionary history of the Western Palearctic blue tits (Cyanistes spp.)–phylogenomic analyses suggest radiation by multiple colonization events and subsequent isolation. Mol. Ecol. 2015, 24(10), 2477–2494.

28. Thompson, J.D.; D.G. Higgins; T.J. Gibson. CLUSTAL W: improving the sensitivity of progressive multiple sequence alignment through sequence weighting, position-specific gap penalties and weight matrix choice. Nucleic Acids Res. 1994, 22(22), 4673–4680.

29. Hall, T.A. BioEdit: a user-friendly biological sequence alignment editor and analysis program for Windows 95/98/NT. in Nucleic acids symposium series. 1999. Oxford.

30. Xia, X. DAMBE5: a comprehensive software package for data analysis in molecular biology and evolution. Molecular biology and evolution. 2013, 30(7), 1720–1728.

31. Kumar, S.; G. Stecher; M. Suleski; M. Sanderford; S. Sharma; K. Tamura. MEGA12: Molecular Evolutionary Genetic Analysis version 12 for adaptive and green computing. Molecular Biology and Evolution. 2024, 41(12), msae263.

32. Lanfear, R.; B. Calcott; S.Y. Ho; S. Guindon. PartitionFinder: combined selection of partitioning schemes and substitution models for phylogenetic analyses. Molecular biology and evolution. 2012, 29(6), 1695–1701.

33. Ronquist, F.; J.P. Huelsenbeck. MrBayes 3: Bayesian phylogenetic inference under mixed models. Bioinformatics. 2003, 19(12), 1572–1574.

34. Edler, D.; J. Klein; A. Antonelli; D. Silvestro. raxmlGUI 2.0: a graphical interface and toolkit for phylogenetic analyses using RAxML. Methods in ecology and evolution. 2021, 12(2), 373–377.

35. Felsenstein, J. Confidence limits on phylogenies: an approach using the bootstrap. evolution. 1985, 39(4), 783–791.

36. Leigh, J.W.; D. Bryant; S. Nakagawa. POPART: full-feature software for haplotype network construction. Methods Ecol. Evol. 2015, 6(9).

37. Librado, P.; J. Rozas. DnaSP v5: a software for comprehensive analysis of DNA polymorphism data. Bioinformatics. 2009, 25(11), 1451–1452.

38. Excoffier, L.; G. Laval; S. Schneider. An integrated software package for population genetics data analysis. Evol. Bioinform. Online. 2005, 1(4), 47–50.

39. Fu, Y.-X. Statistical tests of neutrality of mutations against population growth, hitchhiking and background selection. Genetics. 1997, 147(2), 915–925.

40. Harpending, H. Signature of ancient population growth in a low-resolution mitochondrial DNA mismatch distribution. Hum. Biol. 1994, 591–600.

41. Tajima, F. Statistical method for testing the neutral mutation hypothesis by DNA polymorphism. Genetics. 1989, 123(3), 585–595.

42. Ramos-Onsins, S.E.; J. Rozas. Statistical properties of new neutrality tests against population growth. Mol. biol. evol. 2002, 19(12), 2092–2100.

43. Gracia, E.; J. Ortego; J.A. Godoy; J.M. Pérez-García; G. Blanco; M. del Mar Delgado; V. Penteriani; I. Almodovar; F. Botella; J.A. Sánchez-Zapata. Genetic signatures of demographic changes in an avian top predator during the last century: Bottlenecks and expansions of the Eurasian Eagle Owl in the Iberian Peninsula. PLoS One. 2015, 10(7), e0133954.

44. Aguillon, S.M.; J.W. Fitzpatrick; R. Bowman; S.J. Schoech; A.G. Clark; G. Coop; N. Chen. Deconstructing isolation-by-distance: The genomic consequences of limited dispersal. PLoS genetics. 2017, 13(8), e1006911.

45. Rexer-Huber, K.; A.J. Veale; P. Catry; Y. Cherel; L. Dutoit; Y. Foster; J.C. McEwan; G.C. Parker; R.A. Phillips; P.G. Ryan. Genomics detects population structure within and between ocean basins in a circumpolar seabird: the white-chinned petrel. Molecular Ecology. 2019, 28(20), 4552–4572.

46. Barišić, S.; V. Tutiš; D. Ćiković; J. Kralj; Z. Ružanović. The eagle owl Bubo bubo (Aves: Strigidae) in the Eastern Adriatic (Croatia): the study case of a high-density insular population. Italian journal of zoology. 2016, 83(2), 275–281.

